# Quantifying the contribution of four resistance mechanisms to ciprofloxacin minimum inhibitory concentration in *Escherichia coli*: a systematic review

**DOI:** 10.1101/372086

**Authors:** Boas C.L. van der Putten, Daniel Remondini, Giovanni Pasquini, Victoria A. Janes, Sébastien Matamoros, Constance Schultsz

## Abstract

**Introduction:** Ciprofloxacin resistance in *Escherichia coli* is widespread and adds to the burden of *E. coli* infections. Reviews assessing the genetic basis of ciprofloxacin resistance have mostly been qualitative. However, to allow for the prediction of a resistance phenotype of clinical relevance based on genotypic characteristics, it is essential to quantify the contribution of prevalent genotypic determinants to resistance. We carried out a systematic review to assess the relative contribution of currently known genomic resistance determinants to the minimum inhibitory concentration (MIC) of ciprofloxacin in *E. coli*.

**Methods:** PubMed and Web of Science were searched for English language studies that assessed both ciprofloxacin MIC and the presence or introduction of genetic determinants of ciprofloxacin resistance in *E. coli*. We included experimental and observational studies without time restrictions. Medians and ranges of MIC fold changes were calculated for each resistance determinant and for combinations of determinants.

**Results:** We included 66 studies, describing 604 *E. coli* isolates that carried at least one genetic resistance determinant. Genes coding for targets of ciprofloxacin (*gyrA* and *parC*) are strongest contributors to ciprofloxacin resistance, with median MIC fold increases ranging from 24 (range 4-133) for single Ser83Leu (*gyrA*) mutants to 1533 (range 256-8533) for triple Ser83Leu, Asp87Asn/Gly (*gyrA*) and Ser80Ile/Arg (*parC*) mutants. Other resistance mechanisms, including efflux, physical blocking or enzymatic modification, conferred smaller increases in ciprofloxacin MIC (median MIC fold increases typically around 15, range 1-125). However, the (combined) presence of these other resistance mechanisms further increases resistance with median MIC fold increases of up to 4000, and even in the absence of *gyrA* and *parC* mutations up to 250.

**Conclusion:** This report provides a comprehensive and quantitative overview of the contribution of different genomic determinants to ciprofloxacin resistance in *E. coli*. Additionally, the data demonstrate the complexity of resistance phenotype prediction from genomic data and could serve as a reference point for studies aiming to address ciprofloxacin resistance prediction using genomics, in *E. coli*.

## Introduction

*Escherichia coli* is a Gram-negative bacterium able to adopt a commensal or pathogenic lifestyle in humans and animals.^1^ Adding to the danger of pathogenic *E. coli* is the rise of antimicrobial resistance. *Escherichia coli* has acquired resistance to some of our most important antimicrobials, including aminopenicillins, cephalosporins, aminoglycosides, carbapenems and fluoroquinolones.^2^

Ciprofloxacin is an antimicrobial of the fluoroquinolone class, commonly prescribed for a wide variety of infections including infections caused by *E. coli*.^3^ As is the case for other fluoroquinolones, the substrate of ciprofloxacin is the complex formed by the DNA of the bacterium and either the DNA gyrase enzyme or the topoisomerase IV enzyme.^4–6^ DNA gyrase creates single-stranded breaks in the DNA to negatively supercoil the DNA during replication or transcription.^7^ If ciprofloxacin binds DNA gyrase in complex with DNA, the single stranded DNA breaks cannot be religated and thus accumulate, leading to double stranded DNA breaks.^8^ A similar mechanism is hypothesized for topoisomerase IV.^9^

The mechanisms of ciprofloxacin resistance in *E. coli* have been investigated intensively in the past 30 years. Mutations in genes coding for DNA gyrase and topoisomerase IV contribute to ciprofloxacin resistance in *E. coli*.^10,11^ In addition, efflux pumps may decrease drug accumulation whilst peptides and enzymes may block drug targets or may modify the drug, respectively (Figure 1). Numerous reviews have covered the topic of ciprofloxacin resistance in *E. coli*, but these reviews have been overwhelmingly qualitative in nature.^12–19^

**Figure 1.**
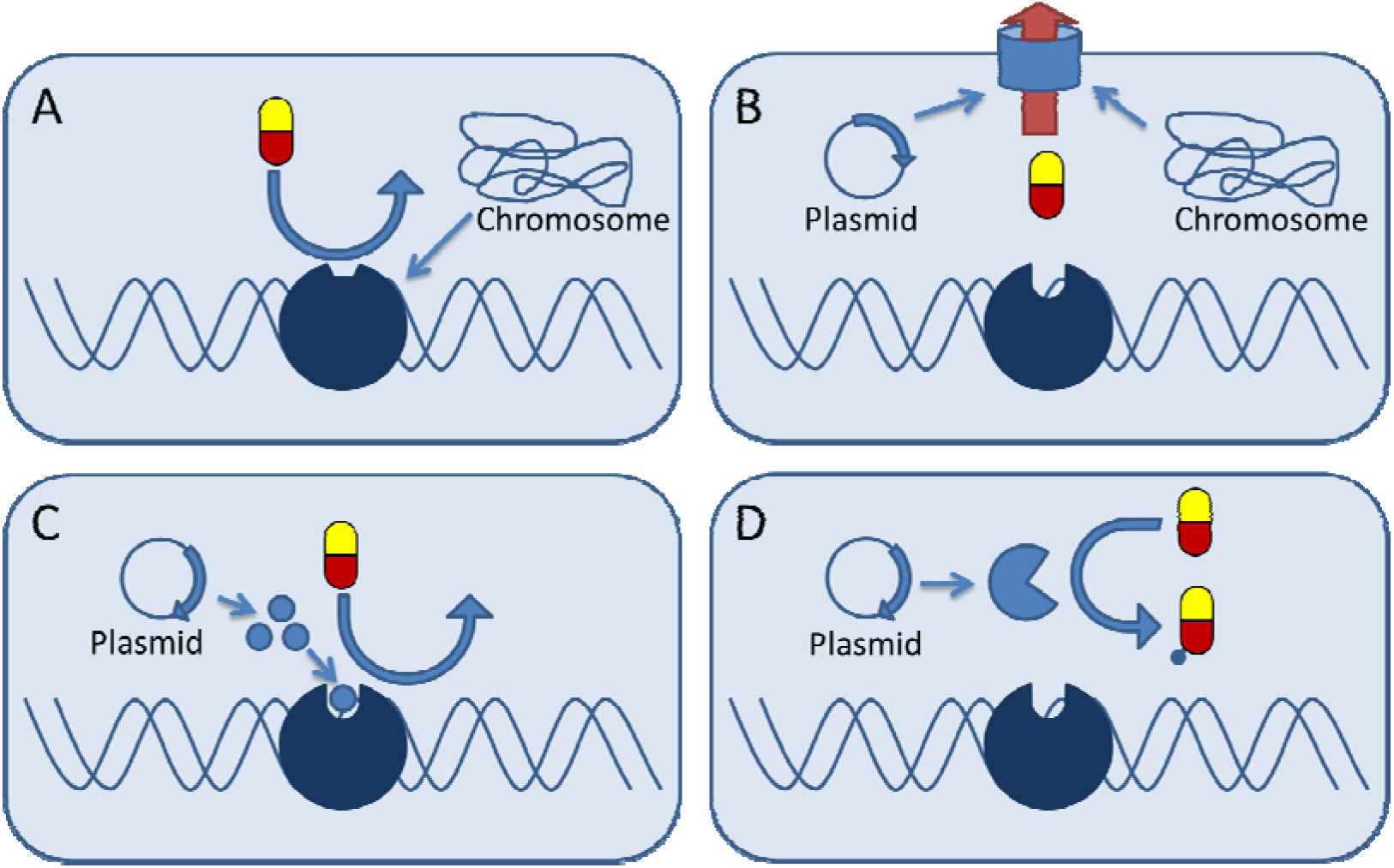
Schematic representation of four mechanisms of ciprofloxacin resistance in *E. coli*. A) Target alteration. B) Decreased ciprofloxacin accumulation. C) Physical blocking of ciprofloxacin target. D) Enzymatic modification of ciprofloxacin.

With the rapidly increasing availability of next generation sequencing technologies, research aimed at the prediction of a resistance phenotype from genomic data is increasing. However, these efforts typically correlate genotypic data to a categorical measure of resistance, while a quantitative resistance phenotype prediction is of clinical relevance. Therefore, we carried out a systematic review, summarizing observational and experimental studies that assessed genetic ciprofloxacin resistance determinants and the ciprofloxacin minimum inhibitory concentration (MIC) conferred by these determinants in *E. coli*, to elucidate how the presence of genomic resistance determinants, either alone or in combination, affects ciprofloxacin MIC in *E. coli*. In addition, we performed an *E. coli* protein network analysis to detect potential additional determinants of ciprofloxacin resistance on the basis of the findings of the systematic review.

## Methods

### Systematic search

The PRISMA 2009 checklist was used as a guide for this systematic review.^20^ PubMed and Web of Science were searched using a defined set of keywords, selecting original research articles in English language reporting on susceptibility test results of *Escherichia coli* isolates measured as Minimum Inhibitory Concentration (MIC) due to genetic modifications identified in clinical, carriage or environmental isolates (observational) or introduced in *E. coli* strains *in vitro* (experimental) (Supplementary methods). No time limits were applied. In addition to the defined search strategy, forward and backward citation searches of reviews and included articles was carried out. The final search was conducted on July 5^th^, 2018.

### Inclusion and exclusion criteria for experimental and observational studies

Articles were not considered eligible for inclusion if they failed to mention any keyword (listed in the supplementary methods) describing ciprofloxacin resistance determinants in title or abstract. Eligible articles were screened by title, abstract and/or full text for inclusion based on the following inclusion and exclusion criteria (Figure 2). Studies could be included as experimental or as observational studies. For inclusion as an experimental study, the study needed to report a ciprofloxacin MIC before and after the introduction of a genetic modification in a single *Escherichia coli* strain. Studies were eligible to be included as observational studies if the ciprofloxacin MIC of at least one *Escherichia coli* isolate was reported, together with the observed genetic determinants of ciprofloxacin resistance. *In vitro* evolution studies where *E. coli* were exposed to ciprofloxacin resulting in decreased susceptibility to ciprofloxacin, were considered observational studies, since mutations are not actively introduced in these studies. Observational studies were excluded if they failed to test for the presence of all of the following resistance determinants: mutations in Ser83 and Asp87 of *gyrA*, mutations in Ser80 and Glu84 of *parC*, mutations in *acrR* and *marR*, presence of *oqxAB*, *qepA*, *qnrA*, *qnrB*, *qnrS* and *aac(6’)Ib-cr*. If studies failed to indicate unambiguously which resistance determinants were tested, the study was excluded.

**Figure 2.**
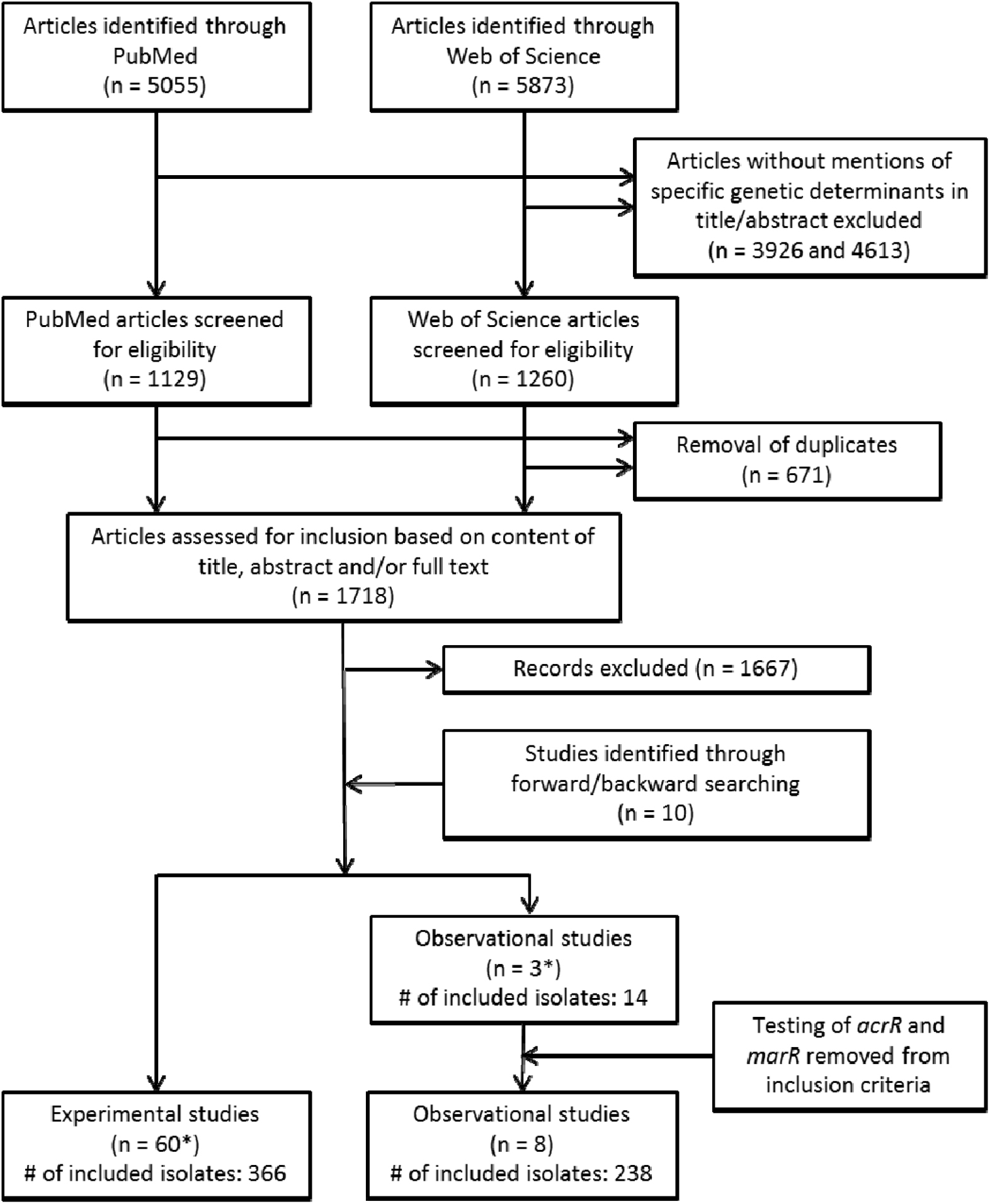
Flow chart adapted from the PRISMA guidelines (Moher 2009), showing the process of including articles starting from a systematic search of PubMed and Web of Science. *2 Studies contributed experimental and observational data, and were thus included for both types of articles.

### Definitions

For this systematic review, the conventional definition of MIC was used, meaning the lowest concentration of ciprofloxacin that inhibits the visible growth of a bacterial culture during overnight incubation.^21^ Clinical breakpoints (≤0.25 mg/L susceptible; 0.5 mg/L intermediately resistant, ≥1 mg/L resistant) and epidemiological cutoffs (0.064 mg/L) were used as defined by EUCAST.^22,23^

A genomic resistance determinant was defined as a mutation in a gene or the presence of a plasmid-mediated gene that decreases ciprofloxacin susceptibility. Since currently four mechanisms of ciprofloxacin resistance in *E. coli* are known, an isolate can possess multiple resistance determinants encoding for multiple resistance mechanisms. In addition, a single resistance mechanism can be encoded by multiple resistance determinants.

Genetic modifications were defined as an experimentally acquired mutation, insertion or deletion of a nucleotide or a sequence of nucleotides in the chromosome. The introduction of plasmid-mediated genes was also considered a genetic modification. Dominance tests as described by Heisig *et al*. were considered experimental evidence.^24^ In short, a dominance test relies on increasing the susceptibility of a bacterium to an antimicrobial, by introducing a plasmid containing the wild type gene that codes for the antimicrobial’s target. In the studies included in this report, the MICs of bacteria with mutations in *gyrA* or *parC* were lowered by introducing a plasmid containing wild type *gyrA* or wild type *parC*.

### Data extraction and analysis

The management of the literature search was performed using Pubreminer (http://hgserver2.amc.nl/cgi-bin/miner/miner2.cgi).

All data on genetic modifications were extracted from the articles or supplementary material, together with MIC data. For experimental data, the MICs of the isolates before and after a targeted genetic modification were extracted to calculate a fold change of ciprofloxacin MIC for each of the *E. coli* isolates.

We calculated how frequently resistance determinants were tested in the experimental data. This frequency is expressed as the number of isolates in which the genetic modification was introduced, divided by the total number of isolates included from experimental studies. The frequency can be used to estimate the strength of evidence per resistance determinant (Table S1). Furthermore, the sample sources, country of origin and isolation date of included *E. coli* isolates were extracted from the observational studies.

The MIC fold change data plot and the correlation matrix were generated using the ggplot2 package RStudio version 1.1.383, running R version 3.4.2. Pearson correlation coefficients were calculated using the stats package and prepared for plotting using the reshape2 package.

### Network construction

To investigate interactions between resistance determinants and to search for potential resistance determinants, a protein-protein interaction network was constructed. The *Escherichia coli* K-12 MG1655 interactome was extracted from the STRING-v10 database.^25^ String-v10 aims to be more complete in terms of coverage of proteins for each organism in comparison to the other meta-interactomes available.^26,27^ The functional association is the basic interaction unit of String in order to link proteins with a functional relation that are likely to contribute to a common biological purpose. Each interaction is derived from multiple sources, and we identify three groups of interactions (Table S3): PI interactions (where at least one physical protein interaction has been tested, imported from primary databases), FP interactions (determined by at least one functional prediction of an algorithm employed by String, genomic information, pathway knowledge, orthology relations) and TM interactions (supported only by automated text-mining of MedLine abstracts and full-text articles). Based on the sources, for each interaction in String a score is calculated, ranging from 0 to 1. In our analysis, only interactions with a score higher than 0.7 were retained (defined as high quality interactions by String), resulting in 3,890 nodes and 32,854 edges (with only 0.06% of the links supported only by TM interactions). Genes resulted by the systematic search were mapped to the EcoGene-3.0 database to obtain *E. coli* K-12 MG1655 identifiers (bnumber)^28^, that were subsequently mapped to the MG1655 interactome.

## Results

### Systematic search

The systematic search yielded 5055 PubMed entries and 5873 Web of Science entries. After removal of duplicates, 1718 unique articles were screened on content by title, abstract and, if necessary, full text. This approach identified 50 articles that were included as experimental studies. Additionally, 10 experimental studies were identified through backward/forward searches in citations of included articles and known reviews. Three articles fulfilled inclusion criteria for observational studies, of which two articles were also included as experimental studies because they provided experimental data as well (figure 2).

The number of *E. coli* isolates which were confirmed to harbour at least one resistance determinant and for which MICs were reported, amounted to a total of 366 isolates from experimental studies (Table S1) and 238 isolates from observational studies (Table S2). A total of 43 different genomic determinants were described in the collected experimental data, of which 21 were shown to have an effect on ciprofloxacin MIC (Table 1).

**Table 1.** Ciprofloxacin resistance mechanisms in *Escherichia coli* and genes involved in these mechanisms. Note that in this overview, only genes are displayed that were shown to have any effect on ciprofloxacin susceptibility when mutations are present (chromosomal genes) or if the resistance gene is present (plasmid-encoded genes).

| Resistance mechanism | Chromosomal genes involved in ciprofloxacin resistance | Plasmid-encoded genes involved in ciprofloxacin resistance |
| --- | --- | --- |
| Target alteration | *gyrA*<sup>12</sup>, *gyrB*<sup>29</sup>, *parC*<sup>11</sup> | - |
| Decreased ciprofloxacin accumulation | *marR*<sup>30</sup>, *acrRAB*<sup>31</sup>, *tolC*<sup>31</sup>,<br>*soxS*<sup>32</sup>, *rpoB*<sup>33</sup> | *qepA*<sup>34</sup>, *oqxAB*<sup>35</sup> |
| Physical blocking of ciprofloxacin target | - | *qnrA*<sup>36</sup>, *qnrB*<sup>37</sup>, *qnrC*<sup>38</sup>, *qnrD*<sup>39</sup>,<br>*qnrE*<sup>40</sup>, *qnrS*<sup>41</sup> |
| Enzymatic modification of ciprofloxacin | - | *aac(6')-Ib-cr*<sup>42</sup><br>*crpP*<sup>43</sup> |

Experimental studies focused primarily on mutations in Ser83 (28% of included isolates) and Asp87 (18%) of *gyrA*, S80 (15%) of *parC* and mutations in *marR* (20%). Of all plasmid-mediated resistance genes, *qnrA* (17%), *qnrS* (12%) and *aac(6’)Ib-cr* (13%) were described most often. The other resistance determinants were tested in less than 10% of the experimentally modified isolates.

### Target alteration mutations in *gyrA, gyrB, parC* and *parE*

Mutations in *gyrA* were the first ciprofloxacin resistance determinants to be discovered (Hooper 1987). Mutations in *parC*, *gyrB* and *parE* were later also proven or implied to decrease ciprofloxacin susceptibility.^11,29,44^ *gyrA* and *parC* mutations that reduce ciprofloxacin susceptibility cluster in regions termed the quinolone resistance-determining regions (QRDRs). Generally, the QRDR of *gyrA* ranges from amino acid Ala67 to Gln106,^45^ and the QRDR of parC from Ala64 to Gln103.^11^ *gyrA* and *parC* mutations accumulate stepwise in *E. coli* when exposed to ciprofloxacin, increasing ciprofloxacin MIC concurrently.^11, 46–48^ The most common initial mutation is Ser83Leu in *gyrA*.^46–48^ In the collected experimental data, this mutation confers a median fold increase in MIC of 24 (range: 4-133x fold increase).^11, 49–55^ This mutation is most often followed by Ser80Ile in *parC*^11,46,48^ and finally by Asp87Asn or Asp87Gly in *gyrA*.^46–48^ As mutations in *gyrA* and *parC* accumulate, ciprofloxacin MIC increases steeply. The ciprofloxacin MIC fold increase for a mutant of Ser83Leu (*gyrA*) and Ser80Ile (*parC*) is 62.5.^51^ A similar double mutant of Ser83Leu (*gyrA*) and Ser80Arg (*parC*) showed a ciprofloxacin MIC fold increase of 125.^53^ For a triple mutant of Ser83Leu, Asp87Asn (*gyrA*) and Ser80Ile (*parC*) the median ciprofloxacin MIC fold increase is 2000.^11,51,54^ A quadruple mutant of Ser83Leu, Asp87Asn (*gyrA*) and Ser80Ile, Glu84Lys (*parC*) has been tested, but this mutant did not show a higher ciprofloxacin MIC than triple mutants within the same study.^11^ In addition, Gly81Asp and Asp82Gly mutations in *gyrA* have been tested. These mutations caused low to no decrease in ciprofloxacin susceptibility (MIC fold changes: 2.6x and 1x, respectively, Table 2).^49,56^

**Table 2.** Medians and ranges of ciprofloxacin MIC fold changes stratified by resistance determinants. Only data from isolates harbouring resistance determinants from a single mechanism are shown.

| Resistance determinant | Median ciprofloxacin MIC fold change (range) | # of isolates | References |
| --- | --- | --- | --- |
| Gly81Asp (*gyrA*) | 2.6 (1-4.2) | 2 | 49,56 |
| Asp82Gly (*gyrA*) | 1 | 1 | 49 |
| Ser83Trp (*gyrA*) | 6.3 | 1 | 10 |
| Ser83Leu (*gyrA*) | 23.8 (4-133.3) | 9 | 11,49–51,53–55 |
| Asp87Asn (*gyrA*) | 15.6 (7.5-15.6) | 3 | 51,54,55 |
| Gly81Asp, Asp82Gly (*gyrA*) | 2 | 1 | 49 |
| Ser83Leu, Asp87Asn (*gyrA*) | 23.8 (15-23.8) | 3 | 51,54,59 |
| Ser83Leu, Asp87Gly (*gyrA*) | 4266.7 | 1 | 77 |
| Asp426Asn (*gyrB*) | 8 | 1 | 29 |
| Ser80Ile (*parC*) | 1 | 1 | 51 |
| Ser83Trp (*gyrA*), Gly78Asp (*parC*) | 33.3 | 1 | 11 |
| Ser83Leu (*gyrA*), Ser80Ile (*parC*) | 62.55 | 1 | 51 |
| Ser83Leu (*gyrA*), Ser80Arg (*parC*) | 125 | 1 | 53 |
| Asp87Asn (*gyrA*), Ser80Ile (*parC*) | 23.8 | 1 | 51 |
| Ser83Leu, Asp87Asn (*gyrA*), Ser80Ile (*parC*) | 2000 (1066.7-2000) | 3 | 11,51,54 |
| Ser83Leu, Asp87Gly (*gyrA*), | 1024 (256-8533.3) | 3 | 11 |
| Ser80Ile (*parC*) |  |  |  |
| Ser83Leu, Asp87Asn (*gyrA*),<br>Ser80Arg (*parC*) | 2258.3 (250-4266.7) | 2 | 11,59 |
| Ser83Leu, D87Y (*gyrA*), Ser80Ile<br>(*parC*) | 256 | 1 | 11 |
| Ser83Leu, Asp87Asn (*gyrA*),<br>Glu84Lys (*parC*) | 533.3 | 1 | 11 |
| Ser83Leu, Asp87Gly (*gyrA*),<br>Glu84Lys (*parC*) | 4266.7 | 1 | 11 |
| Ser83Leu, Asp87Asn (*gyrA*),<br>Ser80Ile, Glu84Gly (*parC*) | 1600 (1066.7-2133.3) | 2 | 11 |
| *acrB*: Gly228Asp | 16.7 | 1 | 61 |
| Δ*acrAB* | 0.1 (0-0.3) | 10 | 30,31,57 |
| Δ*tolC* | 0.3 | 1 | 31 |
| *marR* (various mutations) | 3.5 (1.5-4) | 14 | 60 |
| Δ*marR* | 3.8 (2-218) | 5 | 30,51,54,58,59 |
| *acrR* (various mutations) | 4 (2-16) | 6 | 78 |
| Δ*acrR* | 2.9 | 1 | 51 |
| *soxS*: Ala12Ser | 4 | 1 | 32 |
| *rpoB* (various mutations) | 3 (1.5-3) | 3 | 33 |
| *oqxAB* | 7.5 (2-16) | 17 | 35,62–64 |
| *qepA* | 8.3 (1.9-64) | 13 | 34,52,65–68,79 |
| *qepA*, Δ*marR* | 15 | 1 | 67 |
| *qnrA* (unspecified allele) | 31.3 (20.8-31.7) | 12 | 80 |
| *qnrA1* | 31 (4-66.7) | 37 | 39,50,52,53,81–89 |
| *qnrA3* | 31.3 | 1 | 81 |
| *qnrB1* | 12.5 (4-62.5) | 8 | 52,53,85,87 |
| *qnrB2* | 15.6 (11.8-31.3) | 4 | 81,90 |
| *qnrB4* | 15.6 (15.6-15.6) | 3 | 91 |
| *qnrB5* | 15.6 (15.6-15.6) | 2 | 72 |
| *qnrB6* | 15.6 | 1 | 72 |
| *qnrB19* | 11.9 | 1 | 82 |
| *qnrC1* | 31.3 (15-62.5) | 3 | 59,38,85 |
| *qnrD1* | 15 (7.5-62.5) | 3 | 59,39,85 |
| *qnrE1* | 62.5 | 1 | 40 |
| *qnrS* (unspecified allele) | 12.3 (2-83.3) | 6 | 74,76 |
| *qnrS1* | 33.3 (4-125) | 24 | 39,50,52,53,63,79,81<br>,82,85,87,90,92–94 |
| *qnrS2* | 15 | 1 | 95 |
| *aac(6')Ib-cr* | 6.9 (1-62.5) | 28 | 52,42,71,73–<br>76,79,94,96 |
| *crpP* | 7.5 | 1 | 43 |

Only one *gyrB* mutation (Asp426Asn) was shown to slightly increase ciprofloxacin resistance (Table 2).^29^ We did not find studies that showed a decreased ciprofloxacin susceptibility due to mutations in *parE*. However, a Leu445His mutation in *parE* of *E. coli* caused a 2x fold increase in the MIC of norfloxacin, another fluoroquinolone.^44^

### Efflux pump genes (*acrAB*, *tolC*) and their transcriptional regulators (*marR*, *acrR* and soxS)

As with many other antimicrobials, bacterial efflux pumps also play a role in resistance against ciprofloxacin. Deletion of *acrAB* or *tolC* confers a clear increase in the ciprofloxacin susceptibility of *E. coli* (4-8 fold decrease in MIC).^30,31,57^ Deletions of 14 other genes or operons coding for efflux pumps in *E. coli* did not affect the ciprofloxacin MIC.^31^ The deletion of transcriptional repressors of expression of efflux pumps like *marR* and *acrR* has been shown to affect ciprofloxacin MIC. The only study in our collected experimental data to investigate deletion of *acrR* showed that the MIC tripled after the repressor was deleted.^51^ Nine studies investigated the effects of *marR* deletion or mutation, which reported a median fold increase in ciprofloxacin MIC of 4 (range 1.5-218x fold increase).^30,51,52,54,58–60^ A recent study by Pietsch *et al*. detected mutations in *rpoB* in an *in vitro* evolution experiment.^33^ These mutations arose after accumulation of other mutations, and were shown to increase the ciprofloxacin MIC of a wild type *E. coli* by 1.5-3 fold change (Table 2). The mutations in *rpoB* were shown to increase ciprofloxacin MIC by upregulating the expression of *mdtK* (also known as *ydhE*).

Two experimental studies reported mutations in efflux pump operons, influencing ciprofloxacin MIC. The first mutation was Ala12Ser in *soxS*, leading to higher expression of *acrB*, in turn leading to a ciprofloxacin MIC fold increase of 4.^32^ The second mutation was a Gly288Asp mutation in *acrB* itself, conferring a 16.7 fold increase in ciprofloxacin MIC (Table 2).^61^ This *acrB* mutation however increased susceptibility to other antimicrobials.

### Plasmid-encoded efflux pump genes *oqxAB* and *qepA*

In addition to chromosomally-encoded efflux pumps, the presence of plasmid-encoded efflux pump genes *oqxAB* and *qepA* has been shown to increase ciprofloxacin MIC in *E. coli*.^34,35^ *oqxAB* confers a median fold increase in MIC of 7.5 (range 2-16x fold increase)^35, 62–64^, while *qepA* confers a median fold increase of 4.5 (range 2-31x fold increase, Table 2).^34,52,65–68^

### *qnr* genes

*qnrA* was the first plasmid-mediated quinolone resistance (PMQR) determinant to be discovered.^36^ Qnr proteins are pentapeptide repeat proteins that decrease binding of fluoroquinolones to DNA gyrase by binding the DNA:DNA gyrase complex.^69^ Since 2002, many more *qnr* alleles have been discovered. Currently seven families of *qnr* genes are recognized: *qnrA*, *qnrB*, *qnrC*, *qnrD*, *qnrE*, *qnrS* and *qnrVC*.^70^ In the collected experimental data, all *qnr* families have been tested for their influence on ciprofloxacin MIC of *E. coli*, except for *qnrVC*. *qnr* genes confer ciprofloxacin MIC fold increases between 4 and 125. The median ciprofloxacin MIC fold increase differed per *qnr* allele (Table 2).

### *aac(6’)Ib-cr* and *crpP*

A plasmid mediated mutant *aac(6’)Ib* gene that decreased fluoroquinolone susceptibility in *E. coli* was discovered in 2006.^42^ Until then, *aac(6’)Ib* genes were only known to decrease *E. coli* susceptibility to aminoglycosides. A double mutation in the acetyltransferase-encoding gene enabled the resulting protein to acetylate both aminoglycosides and some fluoroquinolones, including ciprofloxacin. This novel variant, *aac(6’)Ib-cr*, was shown to confer a median fold increase in ciprofloxacin MIC of 6.9 (range: 1-62.5x fold increase, Table 2).^52, 71–76^

The most recently discovered ciprofloxacin resistance determinant in *E. coli* is *crpP*, a plasmid-mediated gene coding for a protein with the putative ability to phosphorylate certain fluoroquinolones such as ciprofloxacin.^43^ *crpP* was first detected in a clinical isolate of *Pseudomonas aeruginosa*, but was shown to confer a 7.5 fold-change increase in ciprofloxacin MIC when conjugated to *E. coli* J53.

### Effect of multiple modifications on MIC

The fold change in MIC of each included experimental isolate was plotted, stratified for the resistance mechanism present (Figure 3). Target alteration resulted in the largest range of MIC fold changes which were on average higher than the fold changes observed as a result of the three other mechanisms. Whilst the presence of determinants representing different ciprofloxacin resistance mechanisms may result in a moderate fold change in MIC, the accumulation of multiple resistance determinants encoding multiple mechanisms of resistance is likely to increase the ciprofloxacin MIC significantly.

**Figure 3.**
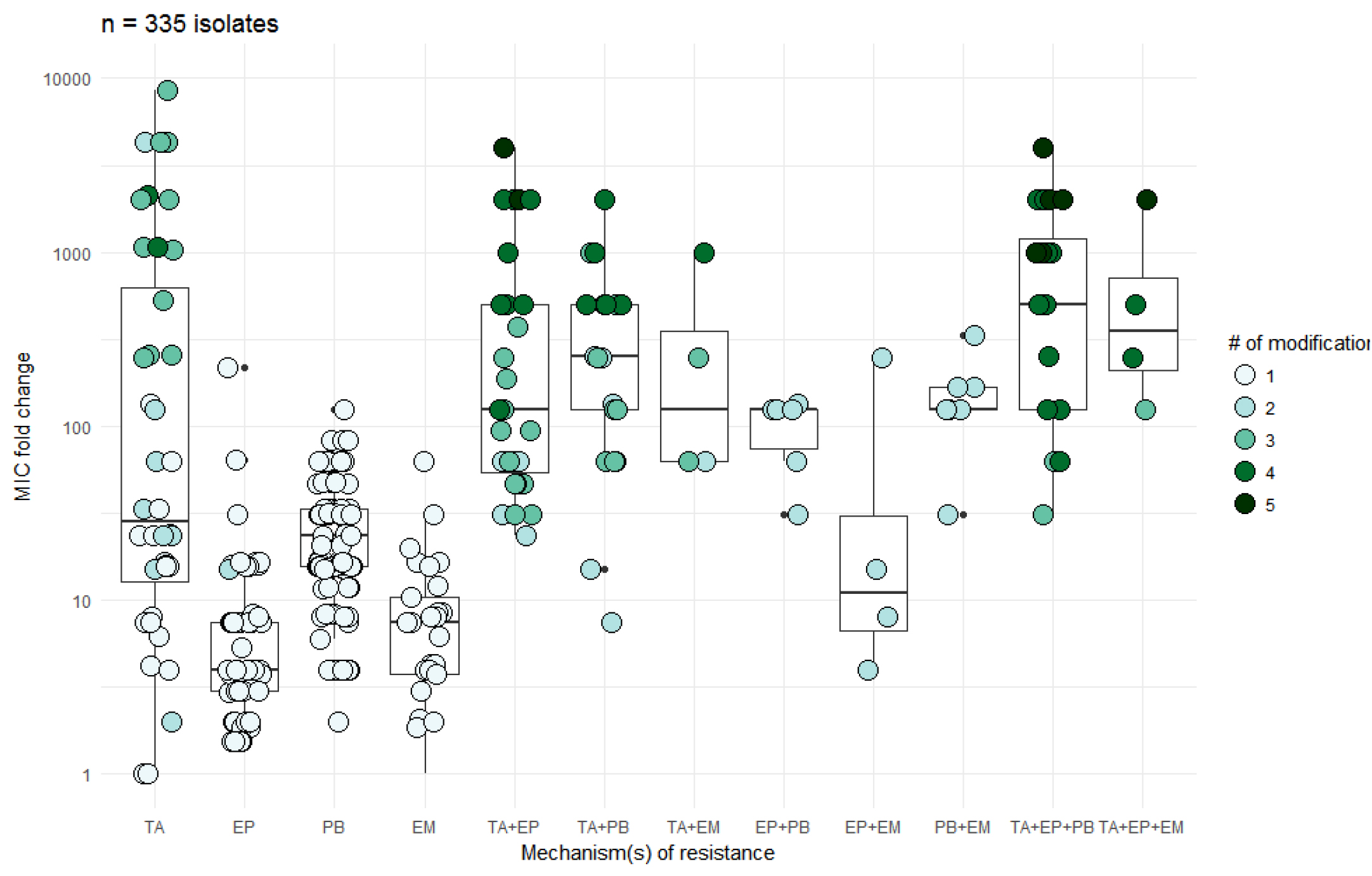
Median fold change (interquartile range) in ciprofloxacin MIC for each resistance mechanism or combination of resistance mechanisms experimentally tested in 366 isolates. Fold changes were calculated by dividing the MIC after modification by the MIC before modification for each isolate. Data points represent single *E. coli* isolates. Darker fill of data points indicates the presence of multiple resistance mutations or resistance genes in the isolate. Isolates that showed a decreased ciprofloxacin MIC after modification (such as deletion of *acrAB* or *tolC*) are not shown but are listed in table S1.^30,31,57^ TA = target alteration (mutations in *gyrA*, *gyrB* or *parC*), EP = efflux pump (mutations in *acrB*, *marR*, *acrR*, *rpoB* or presence of *qepA* or *oqxAB*), PB = physical blocking (presence of *qnrA*, *qnrB*, *qnrC*, *qnrD*, *qnrE* or *qnrS*), EM = enzymatic modification (presence of *aac(6’)Ib-cr* or *crpP*).

### Comparison of experimental and observational data

We compared the findings from the experimental data with susceptibility test results and associated presence of mutations reported for isolates in observational studies. Because studies were excluded if isolates were not tested for the presence of all known resistance encoding determinants, only studies could be included that were published after *oqxAB* was linked to increased ciprofloxacin MIC in 2007.^35^ The description of *crpP* was only recently published and was therefore not used as an inclusion criterion. Only three observational studies reported on the presence of all currently known resistance determinants.^33,97,98^ Since mutations in both *acrR* and *marR* genes were shown to result in no to low fold changes in ciprofloxacin MIC, we added five observational studies that fulfilled all inclusion and exclusion criteria except testing for the presence of mutations in *acrR* and *marR* genes, in a secondary analysis. Thus, eight observational studies published between 2012 and 2018 were included, contributing data on a total of 238 strains (Table S2). The studies reported data on 1 to 92 isolates, with a median of 13.5 isolates per study. Ciprofloxacin MICs of included isolates ranged from 0.015 to 1024 mg/L with a median MIC of 1 mg/L.

We analysed MIC distributions for combinations of resistance determinants that were reported at least five times in the experimental and observational data. These combinations of resistance determinants included the mutation Ser83Leu in *gyrA*, presence of *qnrS1* and presence of *aac(6’)Ib-cr*. Although for most combinations of resistance determinants small numbers of isolates were reported, results of experimental and observational data appear comparable with the exception for the reported MICs for *E. coli* strains solely harbouring *aac(6’)Ib-cr* (Table 3).

**Table 3.** Median ciprofloxacin MICs for three resistance determinants that were reported at least five times in both experimental and observational data. The EUCAST epidemiological cut-off for ciprofloxacin resistance in *E. coli* is 0.064 mg/L.

| Resistance determinant(s) | Median and range of ciprofloxacin MIC in experimental data (mg/L) | Number of isolates in experimental data | Median and range of ciprofloxacin MIC in observational data (mg/L) | Number of isolates in observational data |
| --- | --- | --- | --- | --- |
| Ser83Leu (*gyrA*) | 0.25 (0.06-0.38) | 5 | 0.25 (0.125-64) | 34 |
| *qnrS1* | 0.25 (0.032-1) | 16 | 0.2 (0.1-4) | 19 |
| *aac(6')Ib-cr* | 0.06 (0.004-0.5) | 22 | 0.25 (0.25-0.5) | 5 |

We also examined if certain combinations of resistance mechanisms were more prevalent than others in the observational data. Calculating Pearson correlation coefficients between commonly observed resistance determinants showed that *gyrA* (Ser83, Asp87) and *parC* (Ser80) mutations were positively correlated with each other. Additionally, these three mutations were shown to inversely correlate with the presence of *qnrB* and *qnrS* genes in our observational data. This inverse correlation was not observed with other frequently reported plasmid-mediated resistance determinants such as *aac(6’)Ib-cr* (Figure 4).

**Figure 4.**
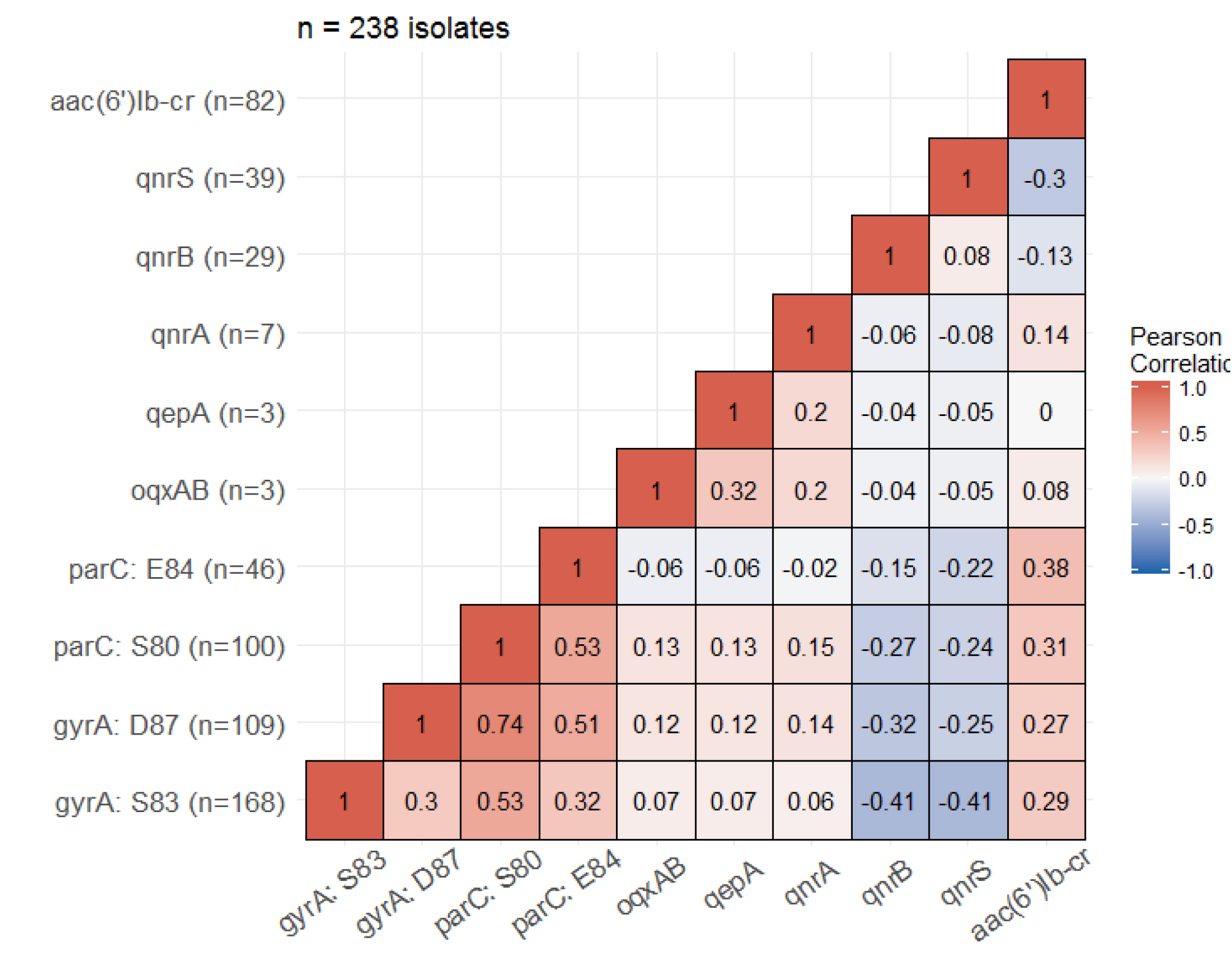
Matrix displaying Pearson correlation coefficients calculated between resistance determinants in a pairwise manner. All 238 strains used for this analysis were screened for all displayed resistance determinants. The reported frequencies of resistance determinants in our dataset are displayed on the y-axis. Full data is provided in table S2.

### Network visualization

In order to get a global picture of the mutation landscape associated with ciprofloxacin resistance, we mapped the selected chromosomal genes onto a Protein-Protein Interaction (PPI) network. The selected genes were evaluated in a wide range of *E. coli* strains, and we mapped them to the String-v10 database referring to the *E. coli* K-12 MG1655 model organism, since it showed the highest number of matching edges and nodes among the strains available in String database. We noted that plasmid-associated genes like *oqxAB* and the *qnr* gene family were not described by interactomes in general, since interactomes mostly describe the core genome. Moreover, some genes (such as *yohG*) could not be mapped because they are not present in *E. coli* K-12 MG1655.

Of the 43 selected genes, 31 (72%) mapped to the PPI network, resulting in a fully connected sub-module. The network highlighted the close relationship between gene connectivity and ciprofloxacin resistance effects: the chosen visualization algorithm showed that genes with similar effects tightly grouped in the interactome (Figure 5). Particularly, the genes that had an increasing effect on ciprofloxacin resistance when mutated seemed to cluster, even if the genes belonged to different resistance mechanisms. As expected, close relationships between particular sets of genes were revealed. Transcriptional regulators such as *marR*, *acrR* and *soxS* were shown to interact with efflux pump genes such as *acrA*, *acrB*, *acrD*, *acrF* and *tolC*. Also, the physical interactions between *gyrA*, *gyrB* and *parC* were depicted in the network.

**Figure 5.**
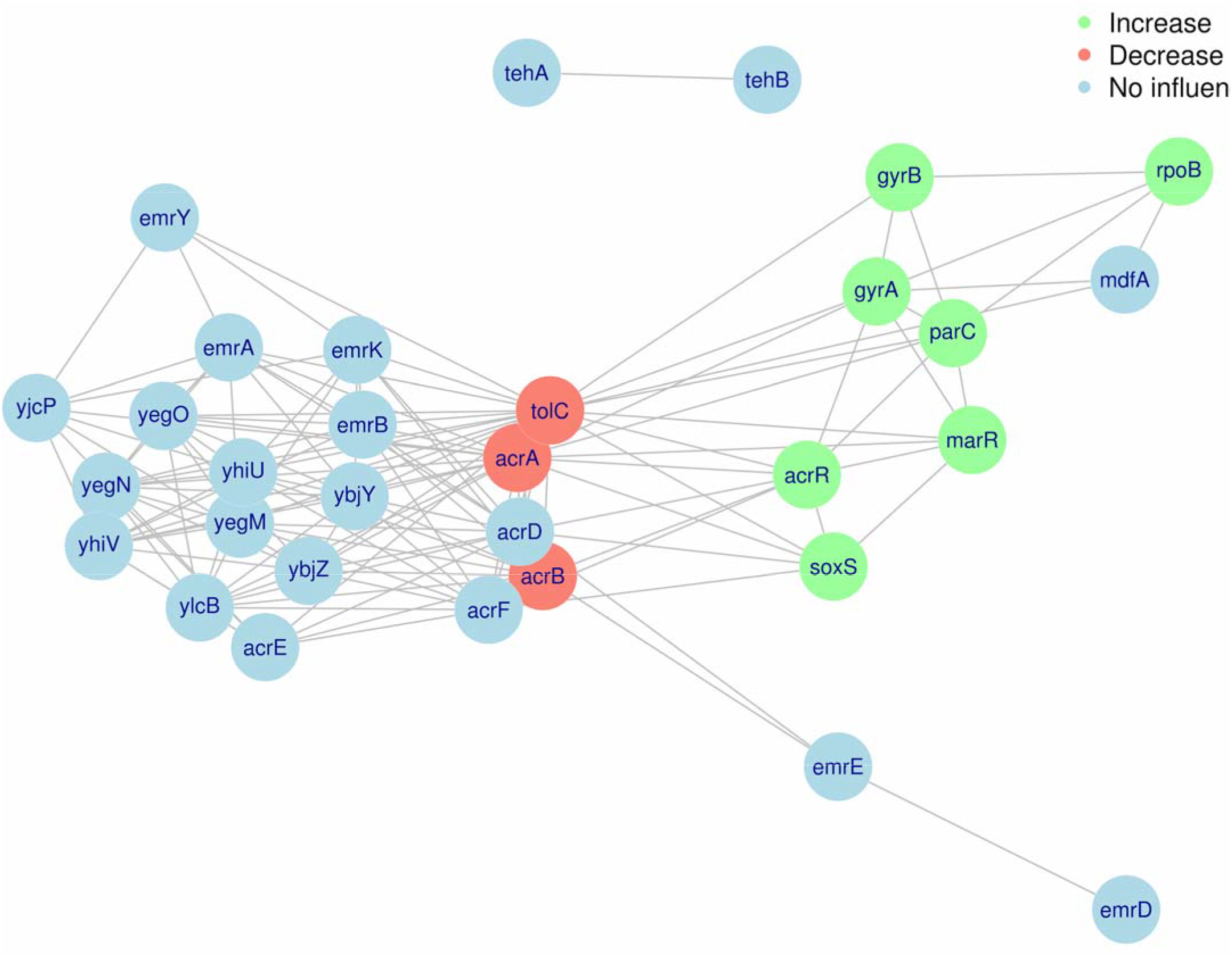
Network of *E. coli* ciprofloxacin resistance-associated chromosomal genes. 31 genes that were examined for their influence on ciprofloxacin and were present in the *E. coli* K-12 MG1655 genome were mapped to the String-v10 PPI database. Genes were coloured green if a mutation conferring increased ciprofloxacin resistance was observed; genes were coloured red when a mutation decreased ciprofloxacin resistance; genes were coloured blue when a mutation showed no effect on ciprofloxacin resistance. The network is displayed by R package iGraph employing the force-directed layout algorithm by Fruchterman and Reingold. The list of edges with corresponding data categories (PI, FP or TM) is available as supplementary table 3.

## Discussion

This report provides a comprehensive and systematic analysis of 66 papers linking genotype of *E. coli* to a quantitative ciprofloxacin resistance phenotype, spanning the years 1989-2018 and amounting to a total of 604 isolates. Ciprofloxacin MIC in *E. coli* is largely affected by target mutations in specific residues in *gyrA* (Ser83 and Asp87) and *parC* (Ser80), conferring median MIC fold increases ranging from 24 for single Ser83Leu (*gyrA*) mutants to 1533 for triple Ser83Leu, Asp87Asn/Gly (*gyrA*) Ser80Ile/Arg (*parC*) mutants. However, accumulation of multiple resistance determinants, including those representing other resistance mechanisms, can increase ciprofloxacin MIC even further, up to MIC fold increases of 4000.

Beside the MIC fold changes that are conferred by resistance determinants, it is important to consider how these genetic resistance determinants are acquired. The SOS response is an important driver of mutation after DNA damage is induced by quinolones such as ciprofloxacin.^99^ Two proteins that are central in the SOS response are LexA and RecA. In the absence of DNA damage, LexA dimers are bound to a SOS box (promoter region of SOS genes) and inhibit expression of SOS genes. If DNA damage is induced, for example through the presence of ciprofloxacin, RecA will bind ssDNA that is a result of the DNA damage. The activated RecA in turn mediates the self-cleavage of LexA, derepressing the SOS box, finally leading to expression of SOS genes and thus the SOS response. This SOS response induces mutations, among others, through DNA damage repair performed by error-prone DNA polymerases.^100^

Currently, four ways are known in which the SOS response affects ciprofloxacin resistance in *E. coli*. First, the SOS response induces a higher mutation rate, making it more likely that ciprofloxacin resistance mutations will arise within a fixed population.^101^ Additionally, if the SOS response is knocked out in *E. coli*, ciprofloxacin MIC decreases. Clinically resistant *E. coli* that had *recA* knocked out showed MIC fold decreases of 4-8.^101^ Furthermore, the SOS response has been shown to induce expression of some *qnr* gene families, for example *qnrB* and *qnrD*.^102,103^ Finally, the SOS response has been shown to promote horizontal transfer of resistance genes when *E. coli* is grown in the presence of ciprofloxacin.^104^

After mutagenesis through mechanisms such as the SOS response, the fitness of the mutant indicates how likely the bacterium is to survive. In absence of ciprofloxacin, *gyrA* mutations and *parC* mutations have been shown to confer limited fitness costs compared to other resistance determinants.^48,51,59,67,75^ Additionally, mutations in *gyrA* and *parC* show positive epistasis, as the MIC fold change of the triple Ser83Leu, Asp87Asn (*gyrA*) and Ser80Ile (*parC*) mutant is higher (2000x fold increase) than would be expected based on the MIC fold changes conferred by the individual mutations (24x, 16x and 1x fold increases, respectively).^51,105^ This epistatic effect thus raises ciprofloxacin MIC very efficiently. This, in combination with the low fitness costs in absence of ciprofloxacin might explain why ciprofloxacin resistance mutations in *gyrA* and *parC* are the most common ciprofloxacin resistance determinants observed in *E. coli*.

Notably, other combinations of resistance determinants also show positive epistatic effects, although the observed effects are weaker. A similar positive epistatic effect was observed for chromosomal *gyrA*/*parC* mutations together with plasmid-mediated resistance determinants *qepA*^67^ and *aac(6’)Ib-cr*.^52,75^ However, experimental studies of combinations of *gyrA* and *parC* mutations with *qnr* genes showed discordant results. One study reported a negative epistatic effect on ciprofloxacin MIC of target alteration mutations with all *qnr* genes tested (*qnrA*, *qnrB*, *qnrC*, *qnrD*, *qnrS*)^59^, and another study observed a similar effect of target alteration mutations with *qnrB*, but the opposite effect for target alteration mutations with *qnrS* in terms of conferred MIC.^52^

The complex relation between *gyrA*/*parC* mutations and *qnr* genes is further illustrated by our findings from the observational data. We observed a clear negative correlation between presence of *gyrA* or *parC* mutations and presence of *qnrB* and *qnrS* genes. This finding is in line with an earlier study that reported an *E. coli* population fixating *gyrA*/*parC* mutations at a reduced rate when the *E. coli* population harboured a *qnr* gene as opposed to when the *E. coli* strain did not harbour a *qnr* gene.^81^ However, no additional fitness costs are usually reported for *E. coli* harbouring both *gyrA*/*parC* mutations and *qnr* genes.^59^ One possible explanation was suggested by the study of Garoff et al., who reported an enhanced fitness cost conferred by *qnr* genes when Lon protease was absent from an *E. coli* genome.^106^ This finding shows that the fitness cost conferred by an antimicrobial resistance gene to an *E. coli* strain can be influenced by genes that do not directly play a role in antimicrobial resistance.

By mapping the selected genes onto a known *E. coli* interactome, we found a clear association between their role in ciprofloxacin resistance and their position in the network, with a significant proximity of genes that produce a similar response in terms of resistance (i.e. increase or decrease). This global picture highlights the presence of common biological functions (mostly associated with the efflux pumps and their regulation), and it suggests that system biology approaches in the future will likely be helpful to identify new targets or specific pathways related to ciprofloxacin resistance or antimicrobial resistance in general. As an example, the position in the network of *acrD* and *acrF* genes, which were not identified as resistance-associated genes in the experiments reported so far, and their biological function as efflux pump protein complexes, suggest that their role in resistance should be more deeply investigated.

Despite its comprehensiveness our study has certain limitations. First, gene expression data are not included in this review because our study aims at prediction of MIC on the basis of a DNA sequence. It has been shown that increased expression of efflux pumps such as *acrAB* or transcriptional regulators of efflux pumps such as *marA* is significantly correlated with increased fluoroquinolone MIC in *E. coli*.^107,108^ Secondly, complex combinations of resistance determinants such as combinations of *gyrA*/*parC* mutations with plasmid-mediated resistance determinants have been reported sparsely in the experimental data. Therefore, the comparison of experimental and observational data for these combinations of resistance determinants is impossible using this dataset. Finally, only currently known ciprofloxacin resistance determinants could be included in this report. The very recent discovery of *crpP* suggests that more resistance determinants or resistance mechanisms are still waiting to be discovered.^43^ Additionally, complex mutation patterns influencing ciprofloxacin resistance through unknown pathways may exist, but current research methods do not usually detect these kinds of effects.

One possible solution for the issues described above would be the use of advanced machine learning algorithms to predict ciprofloxacin resistance. These algorithms should be able to associate large quantities of sequence data with phenotypic metadata in an unbiased manner. One such attempt has been made for ciprofloxacin resistance already.^109^ It was reported that Ser83Phe, Ser83Thr (*gyrA*), Ser80Arg (*parC*) and presence of any *qnr* gene were the most important resistance determinants according to the algorithm used. However, this study used categorical (susceptible or resistant) and not quantitative phenotype data, and included various Enterobacteriaceae species and the results can thus not be directly compared with the data presented here for *E. coli* alone. This is exemplified by the fact that neither Ser83Phe nor Ser83Thr (*gyrA*) were reported in our observational data. For future studies, the data collected for this review could serve as a benchmark, as this review presents a comprehensive set of quantitative data on the contribution of various resistance determinants to ciprofloxacin MIC in *E. coli*.

## Acknowledgments

We wish to thank the COMPARE consortium for support and helpful discussions.

## Transparency

None to declare.

